# Computationally-guided technology platform for on-demand production of diversified therapeutic phage cocktails

**DOI:** 10.1101/2020.01.26.918771

**Authors:** Catherine M. Mageeney, Anupama Sinha, Richard A. Mosesso, Douglas L. Medlin, Britney Y. Lau, Alecia B. Rokes, Todd W. Lane, Steven S. Branda, Kelly P. Williams

**Affiliations:** Systems Biology, Sandia National Laboratories, 7011 East Ave., Livermore, CA, 94550; Energy Nanomaterials, Sandia National Laboratories, 7011 East Ave., Livermore, CA, 94550; Bioengineering & Defense Technologies, Sandia National Laboratories, 7011 East Ave., Livermore, CA, 94550

**Keywords:** Bacteriophage, Phage therapy, genomics

## Abstract

New therapies are necessary to combat increasingly antibiotic-resistant bacterial pathogens. We have developed a technology platform of computational, molecular biology, and microbiology tools which together enable on-demand production of phages that target virtually any given bacterial isolate. Two complementary computational tools that identify and precisely map prophages and other integrative genetic elements (IGEs) in bacterial genomes are used to identify prophage-laden bacteria that are close relatives of the target strain. Phage genomes are engineered to disable lysogeny, through use of long amplicon PCR and Gibson assembly. Finally, the engineered phage genomes are introduced into host bacteria for phage production. As an initial demonstration, we used this approach to produce a phage cocktail against the opportunistic pathogen *Pseudomonas aeruginosa* PAO1. Two prophage-laden *P. aeruginosa* strains closely related to PAO1 were identified, ATCC 39324 and ATCC 27853. Deep sequencing revealed that mitomycin C treatment of these strains induced seven phages that grow on *P. aeruginosa* PAO1. The most diverse five of these were engineered for non-lysogeny by deleting the integrase gene (*int*), which is readily identifiable and typically conveniently located at one end of the prophage. The Δ*int* phages, individually and in cocktails, showed killing of *P. aeruginosa* PAO1 *in vitro* as well as in a waxworm (*Galleria mellonella*) model of infection.

**SIGNIFICANCE STATEMENT:** The antibiotic-resistance crisis in medicine and agriculture has led to renewed interest in phage therapy as an alternative means of treating infection. However, conventional methods for isolating pathogen-specific phage are slow, labor-intensive, and frequently unsuccessful. We have demonstrated that prophages carried by near-neighbor bacteria can serve as starting material for production of engineered phages that kill the target pathogen. Our approach and technology platform offer new opportunity for rapid development of phage therapies against most, if not all, bacterial pathogens, a foundational advance for use of phage in treating infectious disease.

## INTRODUCTION

The emergence of antibiotic-resistant bacteria has become increasingly problematic in recent years. A number of contributing factors have been identified, including overuse of antibiotics in medicine and agriculture, and reduced rate of discovery and production of new antibiotics (1). New treatments for bacterial infections, particularly those caused by antibiotic-resistant pathogens, are needed to combat the crisis. Phage therapy is an attractive alternative to antibiotics that is undergoing a revival (2–4) in the U.S. for this reason.

Bacteriophages are viruses that infect bacterial species, and are regarded as the natural predators of bacteria (5). For use in treating bacterial infections phages hold numerous advantages, including vast natural diversity, low cost, high specificity (such that microbiomes are left intact), and proliferation *in situ* (amplifying and sustaining therapeutic effects) (6). Therapeutic cocktails are generally composed of virulent phages (capable only of the lytic life cycle) isolated from environmental samples. Temperate phages (capable of both lytic and lysogenic life cycles) have typically been excluded because the resulting lysogenic target bacteria would survive to spread resistance to that phage; temperate phages may furthermore carry cargo genes that promote antibiotic resistance and/or bacterial pathogenicity (7, 8). However, there are far more genome sequences available for temperate phages (in the form of prophages integrated within bacterial genome sequences) than for virulent phage genome sequences at NCBI (9), and temperate phages can be converted into non-lysogenic phages using modern genome engineering tools (e.g., by knocking out the integrase gene) (10).

We have developed two complementary computational tools, Islander (11) and TIGER (12), that identify and precisely map integrative genetic elements (IGEs), including prophages, within bacterial (and archaeal) genome sequences. This software reveals the bacterial host, complete sequence, and precise ends of each prophage. Knowledge of the prophage sequence enables its genome engineering to yield a lysogeny-disabled variant safe for therapy, for example through long amplicon PCR and Gibson assembly, with “rebooting” by introduction and growth in the target bacterium (13). Knowledge of the host enables identification of prophage-laden bacteria that are very close relatives of any given target bacterium, such that the phages produced are more likely to be efficacious on the target. This set of computational, molecular biology, and microbiology tools together constitute a new and powerful technology platform for on-demand production of phage therapies against bacterial pathogens (Figure 1). As an initial demonstration of this approach, we produced five engineered lysogeny-disabled phages that kill *P. aeruginosa* PAO1 in liquid culture as well as in a waxworm model of infection. We foresee application of this platform to develop prepared or on-demand phage collections to target nearly any bacterial pathogen, or to control undesirable components of environmental microbiomes.

**Figure. 1.**
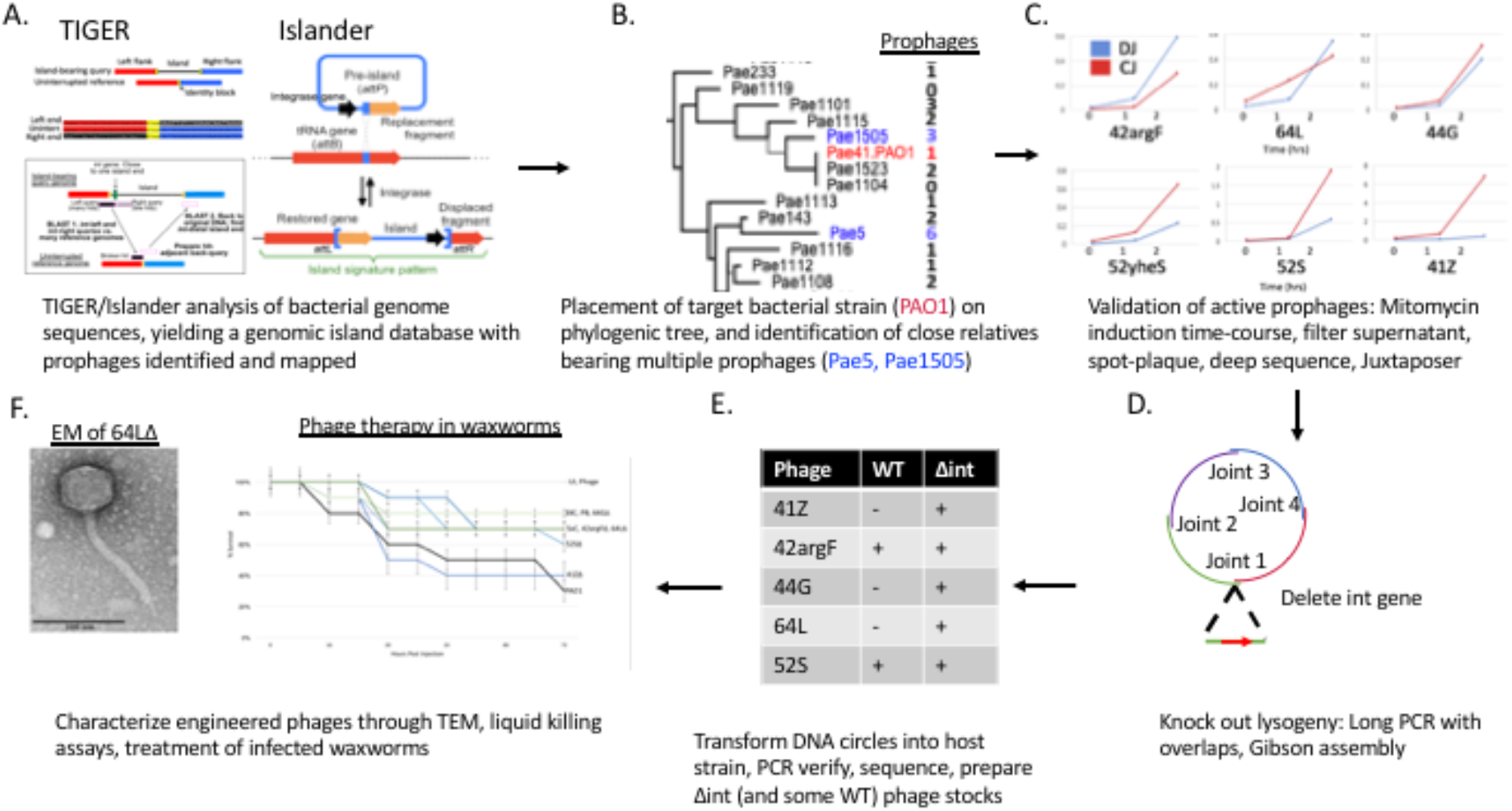
Pipeline for therapeutic phage cocktails. **A)** TIGER and Islander algorithms are run on all sequenced bacterial genomes. Softwares within TIGER and Islander create a database of genomic islands, including prophages. **B)** The target pathogen (PAO1) and close relatives are placed on a phylogenetic tree in the database, such that close relatives bearing multiple prophages can be identified. **C)** Prophage-laden strains are treated with mitomycin C, deep sequenced, and analyzed with Juxtaposer in order to identify prophages capable of mobilization. This step is not essential, but helps ensure that engineering efforts are focused on prophages that can produce active phage. **D)** PCR primers of opposite orientation and with ~40bp overlap are used to generate long PCR amplicons that together span the length of the prophage but do not include its integrase gene. The amplicons are connected up using Gibson assembly, generating a circular Δ*int* phage genome. **E)** The circular Δ*int* phage genome is transformed into a host strain (target pathogen or close relative) for production of Δ*int* phage particles. Plaques are isolated, verified by PCR, and a phage stock prepared (HTL). DNA is then isolated from the stock and sequenced to confirm *int* deletion. **F)** WT and Δ*int* phages are characterized through a variety of methods, including transmission electron microscopy (TEM), *in vitro* killing assays, and treatment of infected *G. mellonella* waxworms.

## RESULTS

### Identification of prophage-laden close relatives of the target bacterial strain

Each IGE encodes an integrase that catalyzes recombination between an attachment site in the circular IGE (*attP*) and one in the bacterial chromosome (*attB*), yielding the IGE integrated into the chromosome flanked by left (*attL*) and right (*attR*) attachment sites. We have developed two complementary algorithms that identify and precisely map *attL* and *attR* for IGEs, including prophages. Islander identifies IGEs with *attB* in a tRNA or tmRNA gene (11) (Figure 1A, right); TIGER identifies IGEs with no bias toward *attB* context (12) (Figure 1A, left). An additional software module identifies prophages among the IGEs (12). Using this approach, we determined the prophage content of the 26 genome-sequenced *Pseudomonas aeruginosa* strains that were available from the American Type Culture Collection (ATCC), totaling 44 prophages. Two *P. aeruginosa* strains (referred to here as Pae5 and Pae1505) were identified that together bear seven potentially useful prophages and are close relatives of our target strain PAO1 (Supp. Figure 1). Pae5 has 12 IGEs, including one filamentous prophages and five additional prophages (41Z, 42argF, 44G, 52yheS, 64L). Pae1505 has ten IGEs, including one filamentous and two additional prophages (43spxA, 52S) (Supp. Table 1). IGE names indicate length (in kbp) and insertion site gene (a single letter representing the identity of a tRNA gene). Filamentous phages tend not to lyse bacteria and are therefore less suitable for therapy (14).

### Ensemble validation of prophages

We sought to determine which of our suspected prophages were able to excise from the bacterial chromosome, produce phage particles, and infect our target strain PAO1. We allowed induction of Pae5 and Pae1505 by treatment for 0, 1 and 2 hours with mitomycin C, and prepared DNA from three sample types: a cell pellet, from a supernatant filtrate (which should contain free phage particles), and a spot plaque of the filtrate grown on a lawn of PAO1. Precise mapping of the prophages by Islander/TIGER allowed design of PCR tests of the circular junction of each excised prophage; PCR products were not detected for the filamentous phages, but were detected for the other seven prophages, in all three sample types of the 2-hr timepoints (Figure 2A). We re-examined all cell pellet DNA samples using deep sequencing, first with our mobilome discovery software Juxtaposer (15) that confirmed the excision products *attP* and *attB*, for six of these seven prophages. We also applied our quantitation tool AttCt (15), to analyze *attP* and *attB* yields (normalized to *attL* and *attR*); higher levels of the former indicate post-excision replication. This analysis reveals (Figure 2B) interesting biological differences in onset of excision and replication rates among the prophages, even along this abbreviated timecourse. One wild-type (WT) phage from each source bacterium, 42argF and 52S, was isolated from each spot plaque by further plaque purification.

**Figure. 2.**
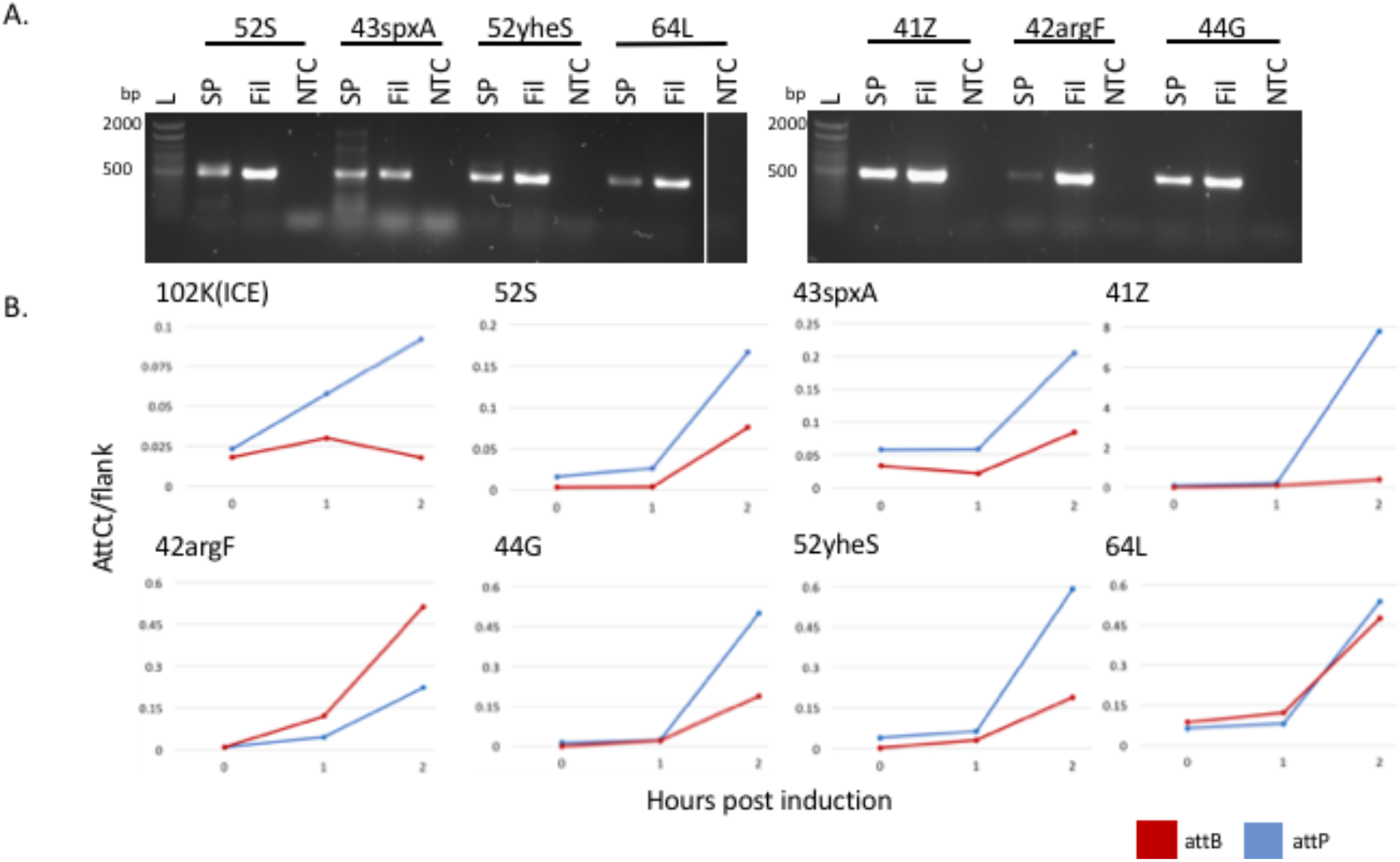
Validation of active prophages. **A)** PCR confirmation of soaked clearings from filtrates spotted onto PAO1 (SP), filtrate from 2 hr MMC induction (Fil), and PB only control (NTC). Primers are listed in Supp. Table 2. All phages are active in both the filtrate and can infect PAO1, shown by the ~500bp band in all SP and Fil lanes. L = Invitrogen 100bp Ladder, 1.2% agarose gel. **B)** AttCt (15) analysis of IGEs identified by TIGER and Islander. Normalized *attP* (blue) and attB (red) counts are shown. Post-excision replication is indicated by elevated levels of *attP* relative to *attB*.

### Genomic content of prophages

Dot plots of the prophage genomes allowed us to reject two (43spxA and 52yheS) as being similar to others, leaving five that we proceeded to develop for therapy (Supp. Figure 2). Annotation of these prophage genomes (Supp. Figure 3) revealed genes for phage structural proteins (capsid, tail, terminase, portal) and promoting lysis (endolysin, holin); only 42argF contained genes predicted to encode deleterious proteins (FimA, a known virulence factor (16); and a *hic* toxin-antitoxin system (17)). 42argF also encodes an endosialidase, which allows phage to recognize and degrade bacterial polysaccharide capsules (18), and an anti-CRISPR AcrF6 protein (Supp. Figure 3).

### Engineering of Δ*int* lysogeny-disabled prophages

Conversion of a temperate phage into a lytic phage requires deletion of at least one key lysogeny determinant. Others have successfully targeted the repressor gene for this purpose (3). For high-throughput genome engineering of phages with diverse bacterial hosts, the repressor gene may be difficult to identify reliably amid other helix-turn-helix proteins. We chose to delete the integrase gene (*int*), which is essential for lysogeny and readily identifiable. Conveniently for deletion, *int* is typically located at one end of the prophage genome (12). The integrase gene and its surrounding regions, including the *attL* and *attR* sites, were deleted by using PCR primers to generate long, partially overlapping amplicons comprising all included prophage sequences, followed by Gibson assembly of the amplicons and transformation into the target host (Figure 1D) (13). Each amplicon was 8-15 kbp in length, producing terminal overlaps with neighboring amplicons of 40-60 bp (Supp. Table 3). Gibson assembly was performed with the 3-5 partially-overlapping amplicons, and the products transformed into electrocompetent (41Z, 42argF, 44G, 52S) or chemically competent (64L) PAO1. The transformed bacteria were plated, and phages recovered from plaques at 16 hours post-plating. PCR was used to verify Δ*int* junctions in the phage genomes. Each phage was propagated on PAO1 in order to create stocks for use in subsequent experiment, and sequenced. No unintended mutations were found for the phages compared to the prophages in the uninduced bacteria, which our re-sequencing did show to bear minor differences to, and to fill gaps from, the published genome sequences.

### Further phage characterization

WT and Δ*int* phages were characterized in further detail (Table 1). No differences in plaque morphology were observed when comparing Δ*int* phages to their parental WT phages (Supp. Fig. 4). Transmission electron microscopy (TEM) was used to determine each phage morphology. All five phages were long-tailed (Siphoviridae). Four of the phages have icosahedral capsids, whereas 52S has a prolate head. Head:tail length ratios were calculated: 41ZΔ*int* = 1:3.4; 42argF WT and Δ*int* = 1:2.9; 44GΔ*int* = 1:2.6; 64LΔ*int* = 1:2.7; 52S = 1:1.3 (Supp. Figure 5). Four of the five phages measured 200-300 nm in length, whereas 52S (both WT and Δ*int*) measured 150 nm in length.

**Table 1.**
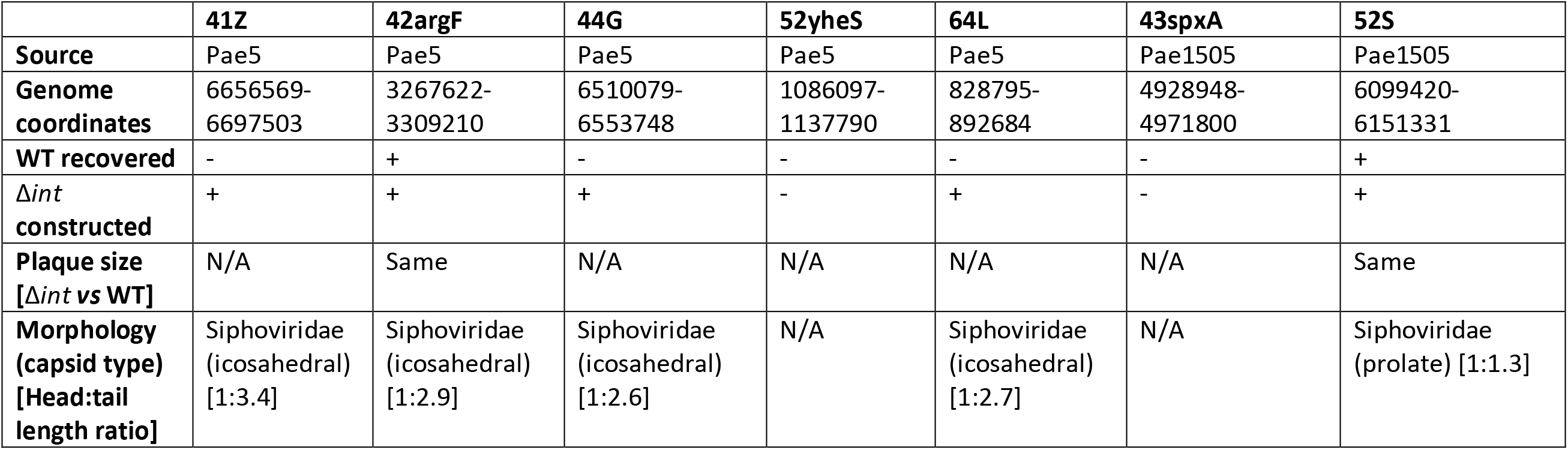
Characterization of phage isolated and engineered from *P. aeruginosa* strains Pae5 (ATCC 2192; NCBI: CH482384.1) and Pae1505 (ATCC 27853; NCBI: CP015117.1).

### Engineered phages kill PA01 in liquid culture

The Δ*int* phages were tested for ability to kill the target pathogen PAO1 in liquid culture. Each phage stock was added to a mid-log phase PAO1 culture at a multiplicity of infection (MOI) of 10 (i.e., 10 phage particles per bacterial cell), and the culture incubated at 37 C with shaking (250 rpm) for 24 hours. Aliquots were removed from the culture at 1-hour intervals for the first four hours following phage addition in order to assess the degree and timing of bacterial death and phage proliferation in the culture. By four hours following phage addition, all cultures exposed to Δ*int* phage showed reduced bacterial titers relative to mock-exposed cultures (i.e., those receiving buffer only), indicating that the each of the Δ*int* phages is capable of killing PAO1 (Figure 3, Supp. Figure 6). Additionally, phage titers increased at least 10-fold, and as much as 1000-fold, in all of the cultures by four hours (Supp. Figure 7).

**Figure 3.**
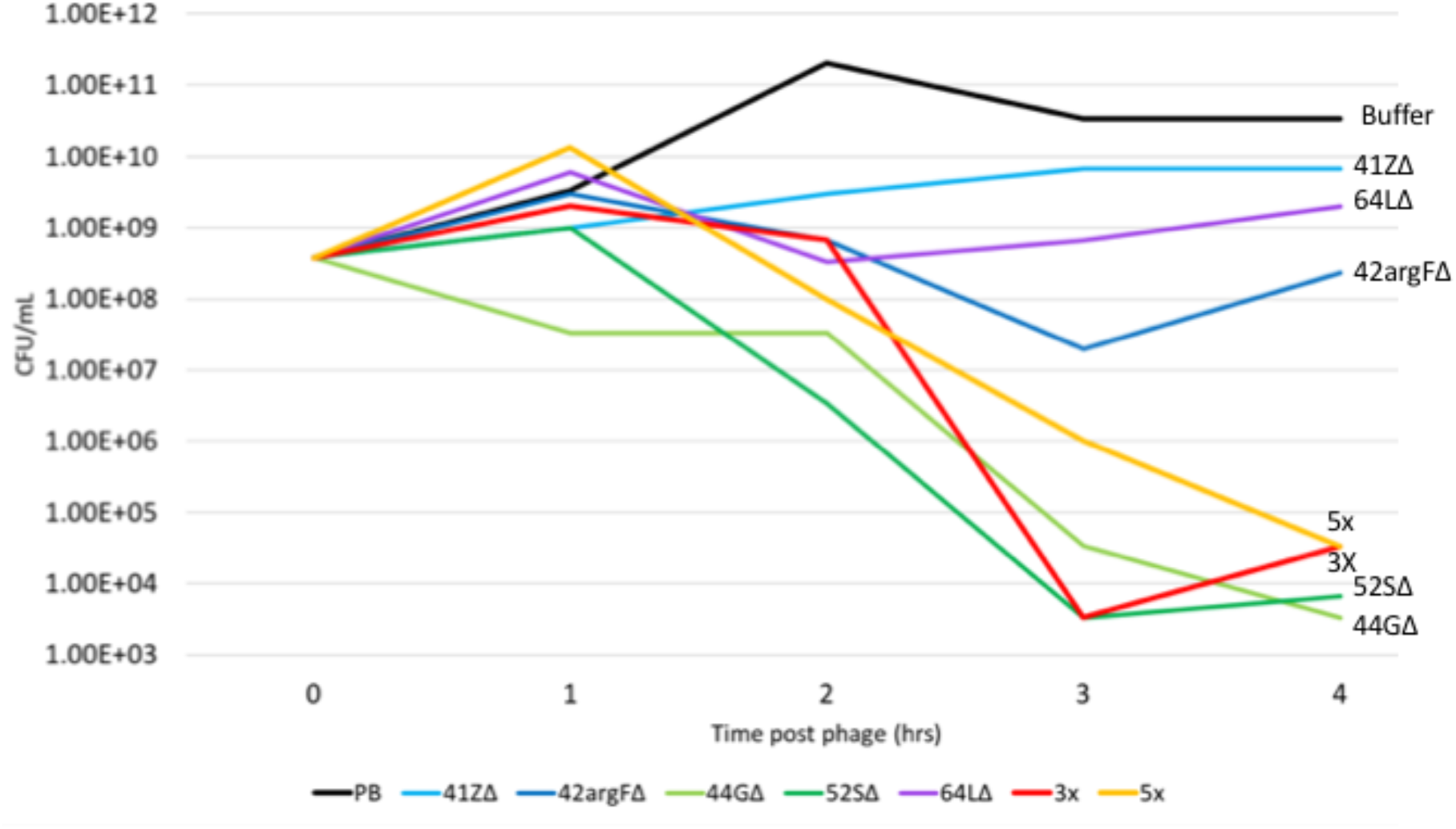
Engineered phages kill PAO1 in liquid culture. PAO1 liquid cultures were grown to mid-log OD_600_=0.4-0.6 then infected with a MOI=10 for each single phage (blue lines), two phage cocktails (green lines) [3X= 44GΔ, 52SΔ, 64LΔ and 5X= 3X plus 41ZΔ and 42argFΔ]or a buffer-only control (black line). Bacterial cells were plated every hour for four hours, and CFU’s were calculated. PAO1 with buffer has increased growth over the time course, while all phage samples show decreased growth over the time course. Decreased growth indicates the phages are killing the PAO1. Replicate number 1 shown, other replicates are shown in Supp. Figures 6.

Two mixtures (cocktails) of Δ*int* phage stocks were designed: cocktail 3X, comprised of phages 44GΔ, 52SΔ, and 64LΔ; and cocktail 5X, comprised of the 3X phages plus 41ZΔ and 42argFΔ. These cocktails were added to PAO1 cultures at an MOI of 10 (MOI of 3.3 for each Δ*int* phage in 3x, and MOI of 2 for each in 5X), aliquots were removed from the cultures at 1-hour intervals over the course of four hours, and the bacterial and phage titers within each aliquot were measured as described above. Both cocktails reduced bacterial titers by more than 10^6^-fold within four hours post-exposure.

### Δ*int* phage therapy protects waxworms from PAO1 infection

We sought to determine whether these phages confer therapeutic effects in the context of a PAO1 infection model. The *Galleria mellonella* larva (waxworm) model is convenient, low cost, and well established for *Pseudomonas* and other bacterial infections and testing of antimicrobial therapies (19) (20–22). Importantly, antimicrobial therapies, including phage therapy, that are efficacious in waxworm models of infection generally also show efficacy in mammalian models of infection (23, 24).

Each larva was injected with 50 CFU of PAO1, and 30 minutes later injected with buffer (negative control), a single Δ*int* phage (MOI of 10), or a phage cocktail (MOI of 10 for sum of phages in cocktail), with each therapy tested in 10 larvae (Figure 4). This experiment was replicated twice more (Supp.Figure 8). Waxworms treated with buffer alone showed only 30% survival by 3 days post-exposure; mortality was due to PAO1 rather than the injection procedure, in that mock-infected larvae (injected with buffer instead of PAO1, and then a second time to simulate treatment) showed 100% survival (gray line). In contrast, larvae treated with single Δ*int* phage showed 40% to 80% survival, the notable exception being 41ZΔ*int*, which had no significant effect on survival (blue lines). Larvae treated with phage cocktails showed further gains in survival, ranging from 70% to 80% (green lines). Treatment with phage in absence of infection had no deleterious effect on survival (gray line). These results indicate that Δ*int* phages, particularly when combined as a cocktail, can protect waxworms against PAO1 infection.

**Figure 4.**
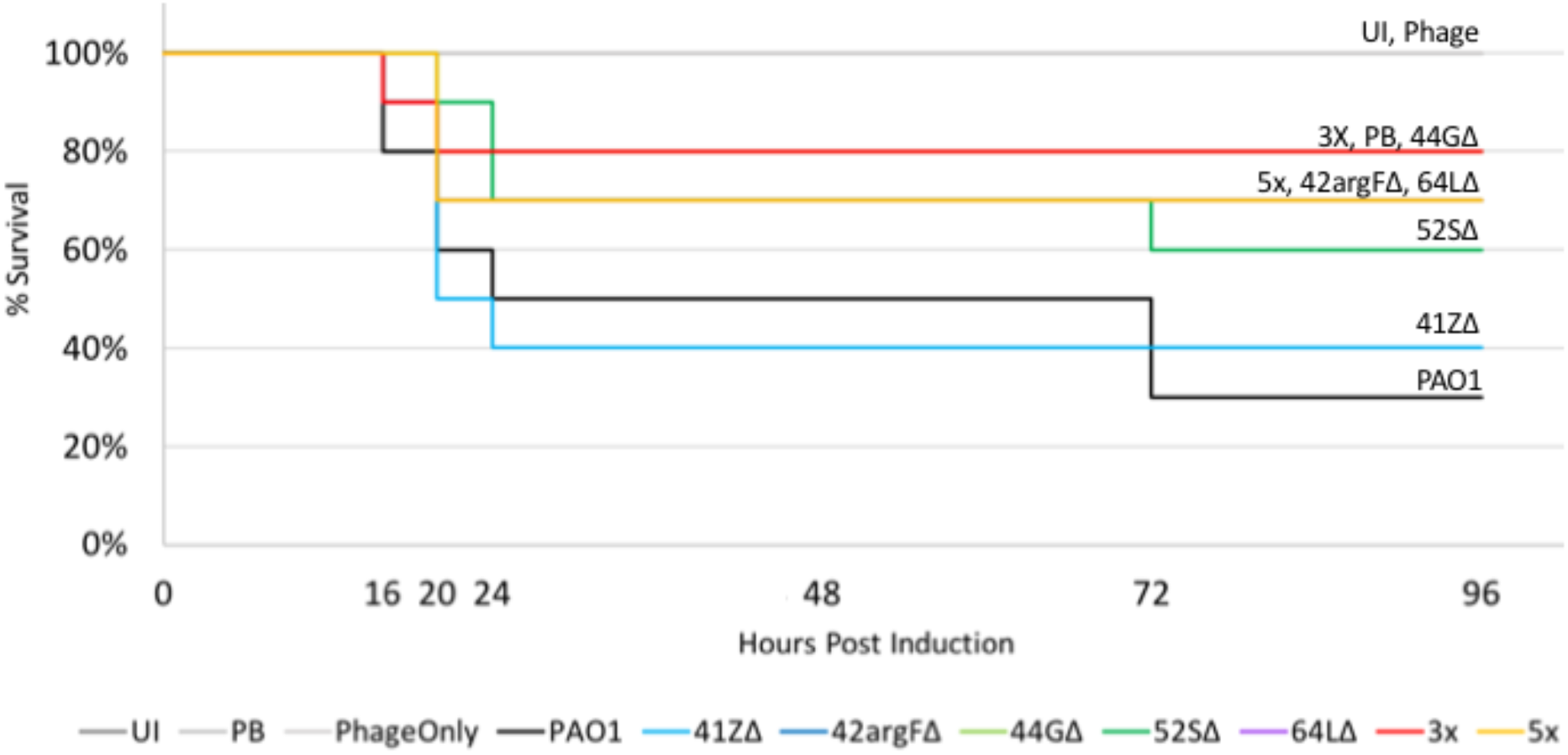
Engineered phages protect *Galleria mellonella* larvae from bacterial infection. Cohorts of ten *G. mellonella* larvae were injected with 10^3^ CFU/mL of PAO1 in the second to last right-proleg, after thirty minutes each larva was injected with a MOI of 10 of either a single phage (blue lines), a phage cocktail (green lines) [3xC = 44GΔ, 52SΔ, 64LΔ and 5xC = 3xC plus 41ZΔ and 42argFΔ], or PB (black line). Three additional controls (grey lines) were used uninjected, PB for both injections, and PB followed by the 5x phage cocktail (PhageOnly). After 72 hours all phages except 41ZΔ saved statistically more larvae than buffer alone. Increased survival indicates the phages are killing the PAO1 in vivo. Replicate one is shown, replicates two and three are shown in Supp. Figure 8.

## DISCUSSION

This report describes an initial demonstration of a new technology platform that enables on-demand production of therapeutic phages against a bacterial pathogen of interest. The first step in our approach is to use two complementary computational tools (Islander, TIGER) to identify and precisely map the prophages present in genome sequences of bacteria closely related to the pathogen. We have carried this out for 2168 genomes (12), and plan to extend our analysis to all available genome sequences. Pre-emptive creation of prophage databases in this manner allows for rapid turn-around in phage therapy cases where time is limited – the physician can use the pathogen’s genome sequence, 16S rRNA gene sequence, or MLST sequences to place it on a phylogenetic tree, to identify close relatives bearing large numbers of prophages. These prophages then serve as starting material for construction of synthetic phages that are engineered for therapeutic use through deletion of genes (such as *int*) essential for lysogeny. The methods described are suitable for high-throughput scale-up.

## MATERIALS AND METHODS

### Prophage detection

Two IGE discovery algorithms, Islander (11) and TIGER (12), were applied to 2023 *Pseudomonas* genomes downloaded from Genbank in October of 2017.

### Phage annotation and comparative genomics

Predicted prophage genomes were annotated. MultiPhATE (25) calling Glimmer (26), Prodigal (27), and Phanotate (28) was used to predict open reading frames. Prokka (29) tFind, and rfind (11) were used for functional annotation. HHPRED (30, 31) was used to further characterize phage functions to compare homology of each protein annotated against the Pfam-A v32.0 and PDB_mmCIF70_28_Nov databases.

Prophage genomes were compared to each other to identify similar prophages. Gepard (32) was used to generate dot plots for the seven non-filamentous prophage sequences. Phage genome maps were created using EasyFig (33).

### Prophage induction

*P. aeruginosa* 2192 (ATCC 39324; Pae5) and *P. aeruginosa* strain Boston 41501 (ATCC 27853; Pae1505) were purchased from ATCC. Strains were grown overnight in LB broth, diluted 1:100 and grown to an OD_600_ of 0.4-0.5. Samples (1 mL) were collected at 0, 1 and 2 hours after treatment with 1 ug/mL mitomycin C. Cells were pelleted at 16000 x g for 2 min; supernatant was removed and filtered through a 0.22 uM filter. Genomic DNA was isolated using the Qiagen DNAeasy Blood and Tissue kit (Qiagen, 69504). Filtrates were spotted onto a lawn of *P. aeruginosa* PAO1 (a kind gift from Annette LaBauve) with 0.5% LB soft agar. Once clearings were observed they were removed and soaked in SM phage buffer (100 mM NaCl, 8 mM MgSO_4_·7H_2_O, 25 mM Tris-HCl pH 7.4) overnight at 4 C and filtered through a 0.22 uM filter.

### Monitoring prophage activity through deep sequencing

*P. aeruginosa* (Pae5 and Pae1505) genomic DNA sequencing libraries were prepared using Nextera DNA library prep kit (Illumina, Cat # FC-121-1031) and utilizing the Nextera DNA Sample Preparation Index Kit (Illumina, FC-121-1011) following the manufacturer’s recommended protocol. DNA samples and libraries were quantified using Qubit high sensitivity DNA assay kit (ThermoFisher Scientific, cat # Q32854). Libraries were pooled in equal quantity and combined to multiplex to make a final library and QC was done again using Qubit quantification kit and Agilent bioanalyzer using high sensitivity DNA chip (Agilent Biotechnology, catalog # 5067-4626). The final combined library was sequenced using Illumina technology on NextSeq 500 sequencer using High output 300-bp Single end Read sequencing kit.

Sequencing reads were processed through BBDuk for quality filtering. The filtered reads were analyzed with Juxtaposer (15) to find mobile elements in the bacterial genomes. Briefly, Juxtaposer searches for recombinant reads relative to the reference genome sequence, which include reads for the *attP* and *attB* products of IGE excision. Finally, attCt software (15) was used to determine counts for each IGE of *attL*, *attR*, *attB*, and *attP*, reporting *attP*/F and *attB*/F values, where F = *attL* + *attR* + 2 *attB*).

### Wild type phage isolation

Soaked filtrates from MMC induction was serially diluted in SM PB. 100uL of *P. aeruginosa* PAO1 was infected with 100uL of each phage dilution, allowed to adsorb 20 min and plated using 4.5mL of 0.5% LB soft agar. Plaques were isolated in SM PB and PCR tests for the circular junction (CJ) were performed to determine which plaque was each phage (Primers listed in Supp. Table 2).

### Gibson assembly of Δ*int* phages

Primers were designed to obtain long PCR fragments with overlapping joints suitable for Gibson assembly strategies (Supp. Table 3). Primers for the integrase deletion were created with artificial 40bp overlaps. Briefly, a 20bp primer was created around the integrase gene and 20bp flanking region was added to the 5’ end of the primer from the opposing end of the Δ*int* circular junction. Long PCR was performed using the NEB Phusion High-Fidelity DNA polymerase master mix. PCR conditions were 98 C for 2 min, (98 C for 30 seconds, Tm-2 for 30 seconds, 72 C for 1 min/kbp) 35 cycles, 72 C for 10 min. NEB Gibson Assembly master mix was used to ligate phage fragments together. 0.3 pmol of each long PCR product was incubated at 50 C for 15 min for 3-fragment phages (41ZΔ, 42argFΔ, 44GΔ), and 60 min for 4 or more fragment phages (52SΔ, 64LΔ).

### Competent cell preparation and transformation

Electrocompetent PAO1 cells were prepared as previously described (34). Briefly, an overnight culture of PAO1 was diluted 1:100 and grown to an OD_600_ of 0.5. Cells were pelted 2 minutes at 16000 x g and supernatant was removed. Cells were washed twice with 300 mM sucrose and resuspended in 300 mM sucrose. For all phages except 64L, 5uL of the undiluted or 1:3 diluted Gibson assembly reaction were delivered into electrocompetent cells using a BioRad Electroporator (2.5 kV, 200 Ω, 25 uF, 2 mm). Cells were allowed to recover in 1 mL of LB media for 1 hr, shaking at 37 C. Cell were pelleted for 2 minutes at 16000 x g and 800 uL of media was removed, cells were resuspended in remaining media. Transformations were diluted in SM phage buffer by 10-fold and 1000-fold. 200 uL of the transformed cells or dilutions were used to infect 100 uL of mid-log PAO, incubating for 20 min, plating with 0.5% LB soft agar and incubating overnight at 37 C.

64L did not produce plaques using electro-transformation protocols. Chemically competent (CC) PAO1 was prepared as previously described (35). Overnight cultures were diluted 1:100 and grown to OD_600_ 0.8. Cells were chilled and harvested by centrifugation, washed with 100 mM MgCl_2_, and incubated on ice in 175 mM CaCl_2_ for 20 min. The final cell pellet was resuspended in 100 mM CaCl_2_. 200 uL of fresh competent cells were incubated with various amounts of Gibson assembly product DNA for 60 min on ice, heat shocked at 42 C for 1 min, chilled and recovered in 1 mL of LB broth for 1 hr prior to plating with 100uL of mid-log PAO1 and 4 mL of 0.5% LB top agar (TA).

### Recovery of Δ*int* mutant phages

Plaques recovered from the transformations were confirmed through PCR with primers designed in the flanking region of the deletion (Supp. Table 2). Once positive plaques were confirmed plaque assays were performed to obtain a web pattern. Web patterns were flooded with 8 mL of SM phage buffer overnight at 4 C, filtered through a 0.22 uM filter. Phage stocks were stored in 0.3% sucrose.

### DNA sequencing

Phage DNA was isolated from stocks with >10^9^ PFU/mL. Phage stocks were treated with 18 U DNase I (New England Biolabs, M0303) and 0.1mg RNase A (ThermoFisher, EN0531) for 1 hour at 37 C followed by 30 min at room temperature. A 2:1 ratio of PEG8000/NaCl_2_ was added and incubated overnight at 4 C. Phenol:chloroform extraction was performed. Two volumes of 95% EtOH and 1/10 volume of 3 M sodium acetate was added to the final aqueous layer and allowed to incubate at room temperature for 30 min. DNA was spooled onto plastic loops and placed in 70% EtOH, spun and resuspended in TE.

Phage sequencing libraries were prepared using Nextera DNA Flex Library prep kit (Illumina, 20018705) and utilizing the Nextera DNA CD Index Kit (Illumina, 20018708) following the manufacturer recommended protocol. DNA samples and libraries were quantified using Qubit high sensitivity DNA assay kit (ThermoFisher Scientific, Q32854). Libraries were pooled in equal quantity and combined to multiplex to make a final library and QC was done again using Qubit quantification kit and Agilent bioanalyzer using high sensitivity DNA chip (Agilent Biotechnology, 5067-4626). The final combined library was sequenced using Illumina technology on NextSeq 500 sequencer using mid output 150-bp paired-end (PE) sequencing kit (Illumina, 20024905).

### Electron microscopy

Electron microscopy grids were prepared with fresh lysates at >10^9^ pfu/mL. 10uL of lysate was added to a carbon grid (Ted-Pella, 1813) and allowed to incubate 10 min. The grids were washed twice with 10uL of water for 2 min. Grids were stained with 10uL of uranyl acetate alternative stain (Ted Pella, 19485, using Gadolinium Acetate Tetrahydrate) for 2 min and wicked off. Grids were allowed to dry at RT in a chemical fume hood for 1 hour and stored in a grid box until imaging.

Transmission electron microscopy observations were conducted using a Themis Z Transmission Electron microscope (Thermo Fisher Scientific, Hillsboro, Oregon, USA) operated in scanning transmission electron microscopy (STEM) mode at an accelerating voltage of 300 kV. Image signals were collected using a high angle annular dark field (HAADF) detector, which is sensitive to the heavy elements (Gd) employed in the stain. In the micrographs presented here, the dark-field image contrast has been inverted, with the heavy-metal stain showing as dark regions on the images.

Immediately prior to observation, specimens were plasma cleaned (Mobile Cubic Asher, IBSS Group, Burlingame CA, USA) to reduce the build-up contamination under exposure to the focused electron beam. Plasma cleaning was conducted for 5 minutes using ambient air at a base pressure of 10^−4^ torr and power setting of 34 W.

### Liquid killing Assays

Overnight PAO1 cultures were diluted 1:100 to an OD_600_ of 0.02-0.04 in LB broth. Cultures were grown to an OD_600_ 0.4-0.5. Cells were pelleted by centrifugation 3000 RPM for 5 min. Bacterial pellets were resuspended in 1 mL PB containing a single phage stock at MOI=10 or a phage cocktail with a total MOI=10; 5 mL of LB broth was added to each culture. Cultures were grown for 24 hr, removing 0.5 mL samples at 0, 1, 2, 3, 4, and 24 hours after phage addition. Each sample was processed for PFU, CFU and OD_600_ measurements.

Bacterial cell counts were performed by serially diluting the sample in LB broth to 10^−8^ and spotting 3 uL of each dilution onto a solid LB agar plate and incubating 37 C for 16 hr. PFUs were determined by centrifugation of each sample at 3000 RPM for 5 min, removing 100 uL of supernatant, serial diluting the supernatant to 10^−10^, followed by spotting 3 uL of each dilution onto a 0.5% soft agar overlay plate. Bacterial and phage titers were calculated by counting the number of colonies or plaques, respectively, followed by conversion to CFU/mL or PFU/mL.

### Waxworm phage therapy

*Galleria mellonella* larvae were purchased from Timberline (Marion, IL) and stored at 15 C until use within two weeks. Only larvae weighing between 0.25-0.35 g were used for injection experiments. Larvae were equilibrated at room temperature 4 hr prior to injection. 5uL of PAO1 (10^3^ CFU/mL) was injected into the second to last right-proleg using a 250 uL, model 1725 LT syringe (Hamilton, 81101) and PB600 repeating dispenser (Hamilton, 83700). 30 min after the PAO1 injection, a 5 uL phage injection of MOI=10 (10^4^ PFU/mL) was injected into the second to last left proleg, and incubated at 37 C for 72 hr. Death was assessed by melanization and lack of movement at 16, 20, 24, 48, 72 hours post injection. Kaplan-Meier survival curves were calculated as described (36).

## ACKNOWLEDGMENTS

This work was supported by the Laboratory Directed Research and Development program at Sandia National Laboratories, which is a multimission laboratory managed and operated by National Technology and Engineering Solutions of Sandia LLC, a wholly owned subsidiary of Honeywell International Inc. for the U.S. Department of Energy’s National Nuclear Security Administration under contract DE-NA0003525.

We thank Anette LaBauve for kindly donating the *P. aeruginosa* PAO1 strain.

