## Supplemental Figures and Tables for "Computationally-guided technology platform for on-demand production of diversified therapeutic phage cocktails"

**SUPPLEMENTAL INFORMATION**

**Supplemental Figures**

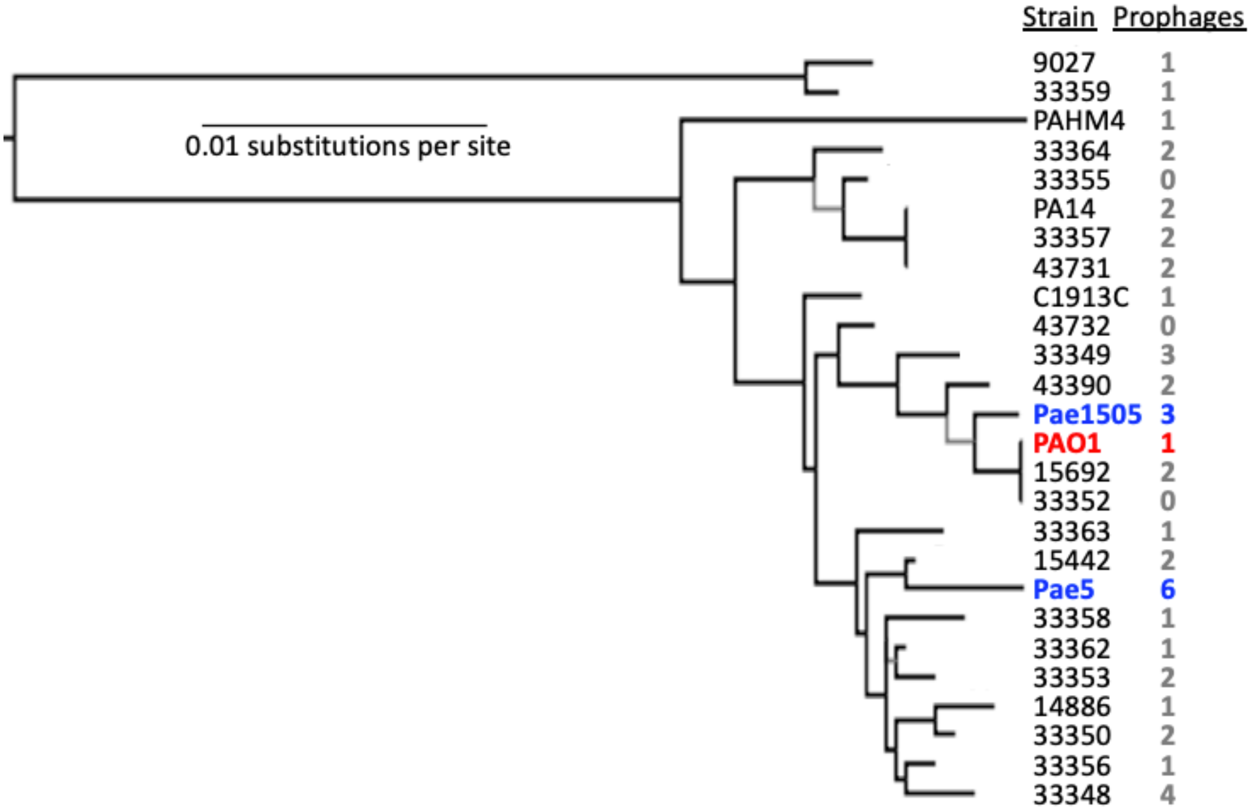


**Supplemental Figure 1. Phylogenetic tree and prophage counts of *Pseudomonas aeruginosa* strains available from ATCC**. For the 25 genome-sequenced *Pseudomonas aeruginosa* strains available from ATCC in October, 2017 (and *P. aeruginosa* PA14, and as an outgroup to root the tree but not shown, *Pseudomonas pseudoalcaligenes* KF707), the nucleotide sequences of the seven genes used for multi-locus sequence typing (37) were taken. Sequences were aligned using Muscle separately for each gene, and the alignments were concatenated, yielding a 2915 bp alignment. A maximum likelihood tree was built using FastTree 2.1.10. Nodes with support values below 0.5 are shown in gray. Strains are designated by ATCC number or other names, with those used in this study in color; the number of prophages determined by Islander/TIGER (12, 38) are given.

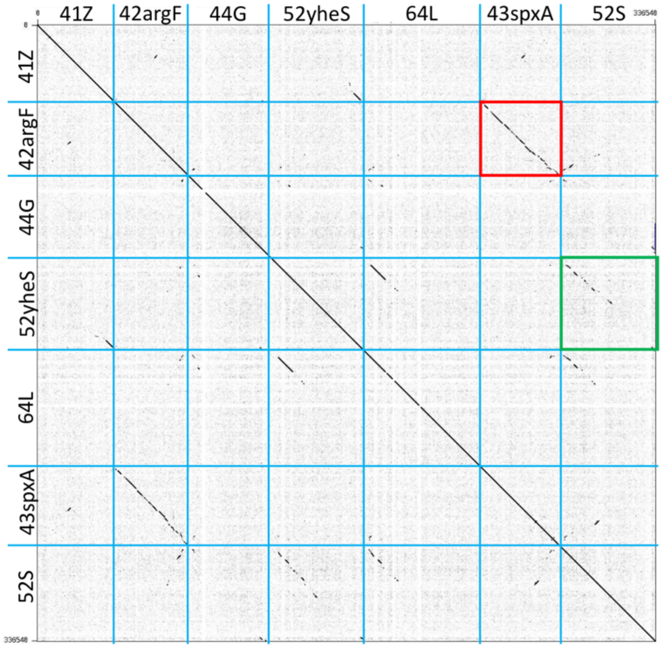


**Supplemental Figure 2. Genomic dot plot of seven active prophages.** The genomes of all seven phages were concatenated into a single file, applying Gepard software (32) to create a dot plot. Phage names are listed on the axes. The diagonal line has a few gaps that result from gaps in the *P. aeruginosa* genome sequencing. The red box shows the high level of similarity between 43spxA and 42argF. The green box shows similarity between 52S and 52yheS.

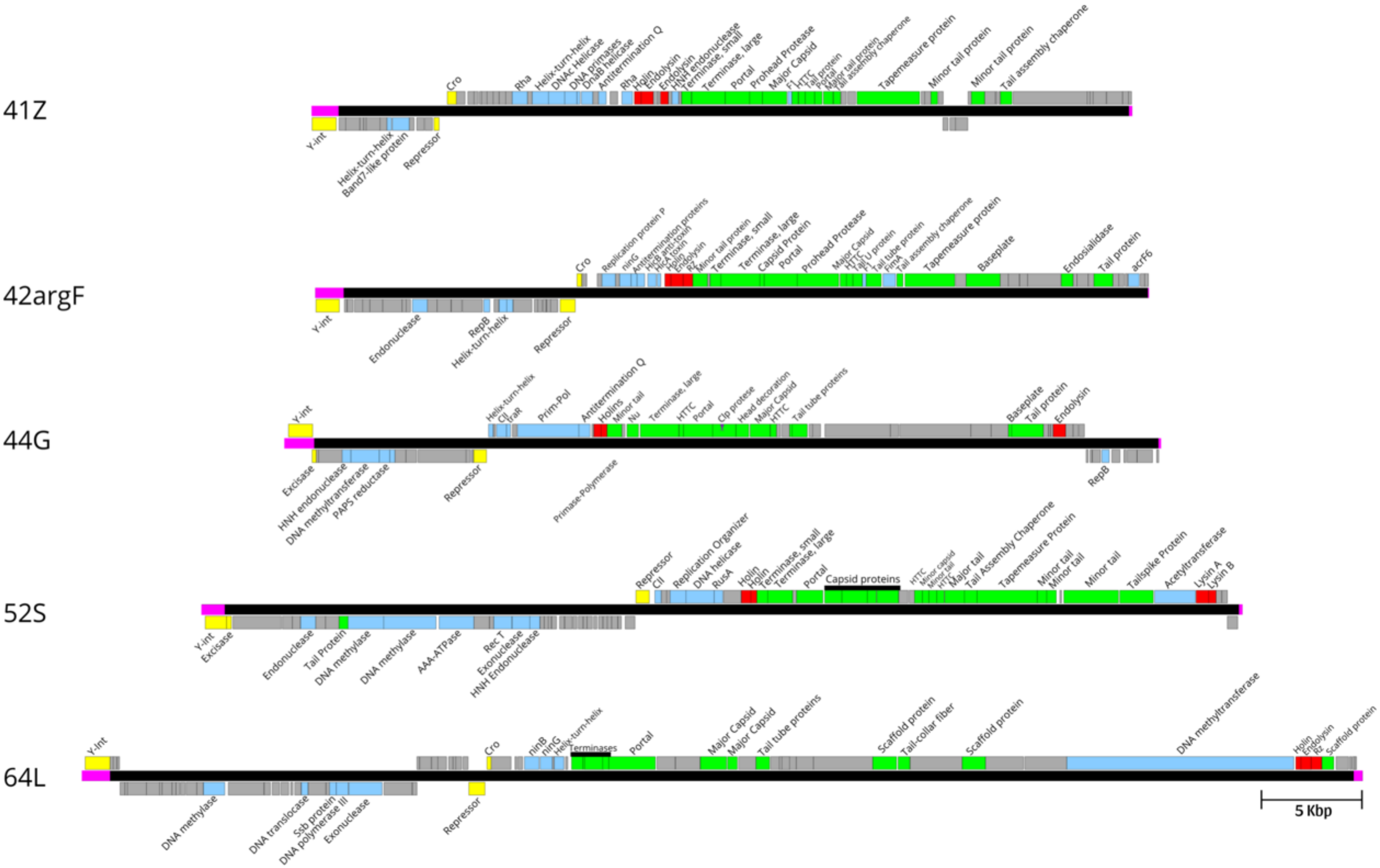
41Z
Cro
Rha
Helix-turn-helix
DNAc Helicase
DNA primases
DnaB helicase
Antitermination Q
Rha
Holin
Endolysin
HNH endonuclease
Terminase, small
Terminase, large
Portal
Prohead Protease
Major Capsid
HTTC
Tail protein
Tapemeasure protein
Minor tail protein
Minor tail protein
Tail assembly chaperone
Y-int
Helix-turn-helix
Band7-like protein
Repressor
42argF
Cro
Replication protein P
ninG
Antitermination proteins
Holin
Endolysin
Rz
Minor tail protein
Terminase, small
Terminase, large
Capsid Protein
Portal
Prohead Protease
Major Capsid
HTTC
Tail tube protein
Tail assembly chaperone
Tapemeasure protein
Baseplate
Endosialidase
Tail protein
acrF6
Y-int
Endonuclease
RepB
Helix-turn-helix
Repressor
44G
Y-int
Helix-turn-helix
CII
Prim-Pol
Antitermination Q
Holins
Minor tail
Nu
Terminase, large
HTTC
Portal
Clp protese
Head decoration
Major Capsid
HTTC
Tail tube proteins
Baseplate
Tail protein
Endolysin
Excisase
HNH endonuclease
DNA methyltransferase
PAPS reductase
Repressor
Primase-Polymerase
RepB
52S
Repressor
CII
Replication Organizer
DNA helicase
RusA
Holin
Holin
Terminase, small
Terminase, large
Portal
Capsid proteins
HTTC
Major tail
Tail Assembly Chaperone
Tapemeasure Protein
Minor tail
Minor tail
Minor tail
Tailspike Protein
Acetyltransferase
Lysin A
Lysin B
Y-int
Excisase
Endonuclease
Tail Protein
DNA methylase
DNA methylase
AAA-ATPase
Rec T
Exonuclease
HNH Endonuclease
64L
Y-Int
Cro
ninB
ninG
Helix-turn-helix
Terminases
Portal
Major Capsid
Major Capsid
Tail tube proteins
Scaffold protein
Tail-collar fiber
Scaffold protein
DNA methyltransferase
Holin
Endolysin
Rz
Scaffold protein
DNA methylase
DNA translocase
Ssb protein
DNA polymerase III
Exonuclease
Repressor
5 Kbp

**Supplemental Figure 3. Genomic maps of the five prophages selected for engineering.** Genes are marked above the line for forward orientation, and below for reverse orientation. Gene functions were predicted using HHpred (30, 31) searching the Pfam-A and PDB protein databases. Predicted functional categories were: lysogeny genes (yellow), lytic genes (red), structural genes (green), functional genes in other categories (blue) and hypothetical genes (grey). Abbreviations: Y-int, tyrosine integrase; HTTC, head-to-tail connector protein; ssb, single-stranded DNA binding protein. The pink boxes represent genome segments were deleted during Δ*int* phage production. The scale bar represents 5 kbp.

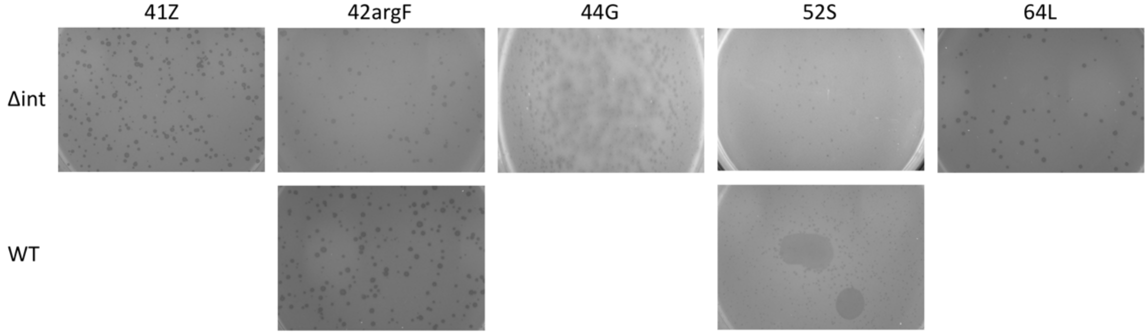


**Supplemental Figure 4. Plaque Morphology for each WT and Δ*int* Phage**
Plaque Assays were performed and plates with 50-100 plaques were imaged after 24 hours of incubation.

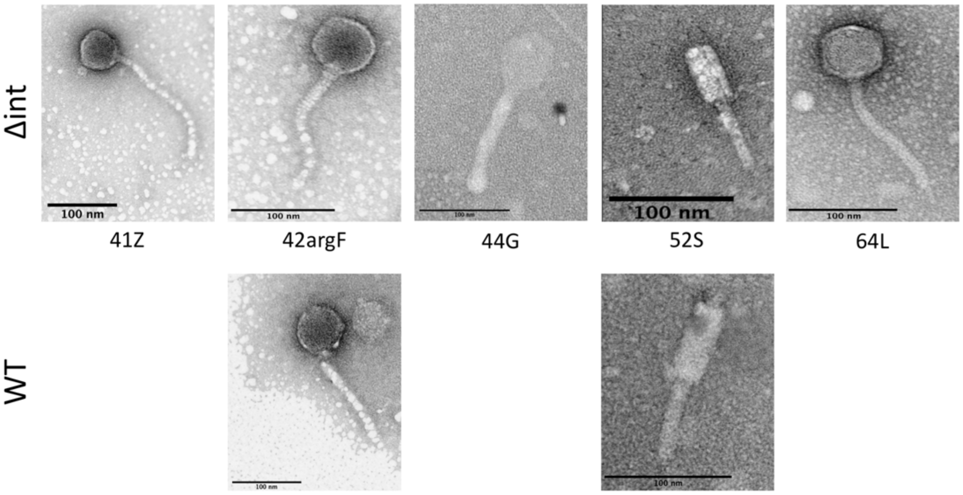


**Supplemental Figure 5. Electron microscopy (EM) of phages.** EM grids were prepared with $>10^9$ pfu/mL of fresh lysate, stained with uranyl acetate alternative stain (Ted Pella, 19485, using Gadolinium Acetate Tetrahydrate) , and imaged with a Themis Z transmission electron microscope operated in HAADF-STEM mode (image contrast inverted for clarity). Scale bars are shown, representing 100 nm.

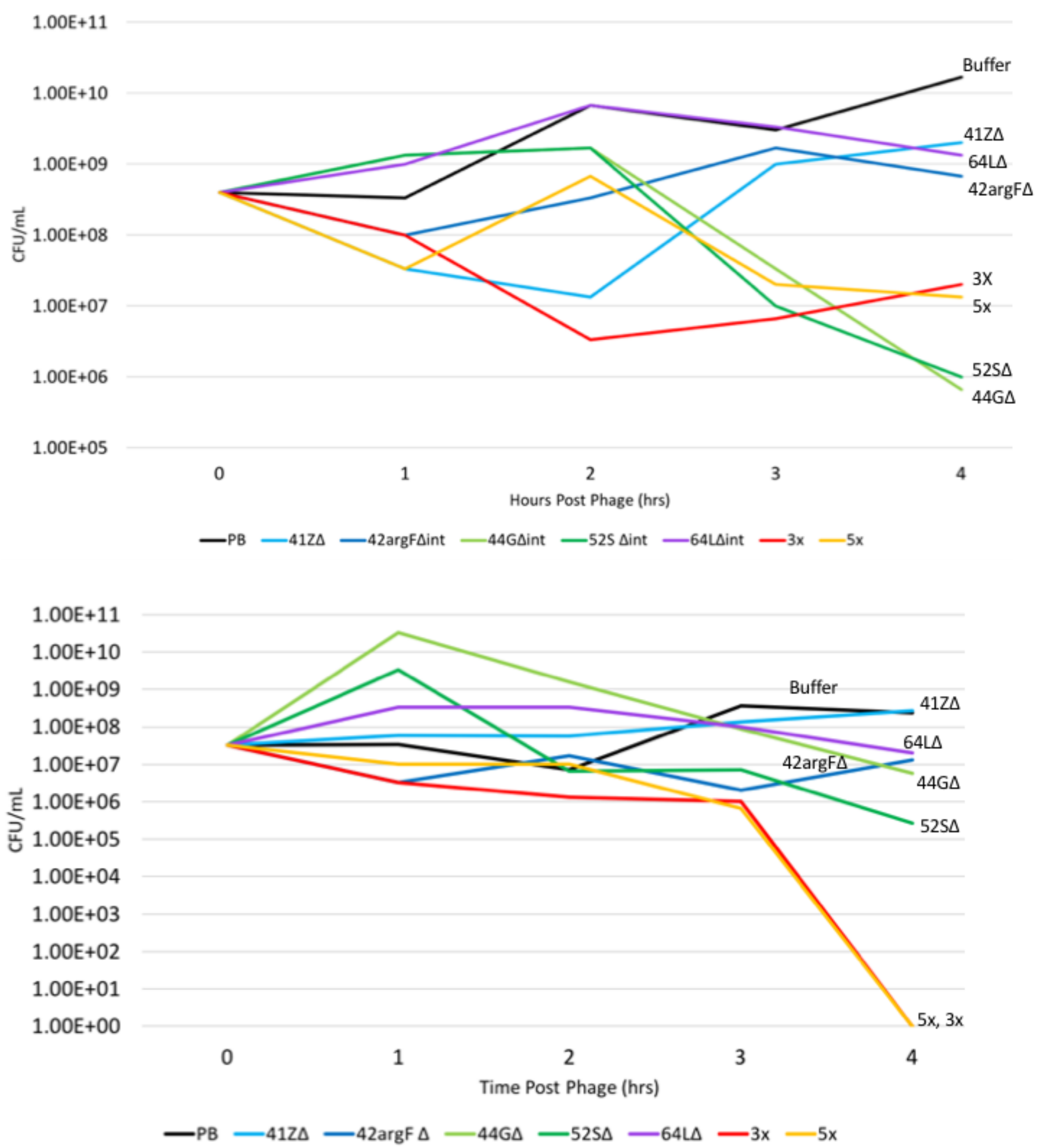


**Supplemental Figure 6. Engineered phages kill PAO1 in liquid culture.** Two additional independent replicates of the experiment in Figure 3.

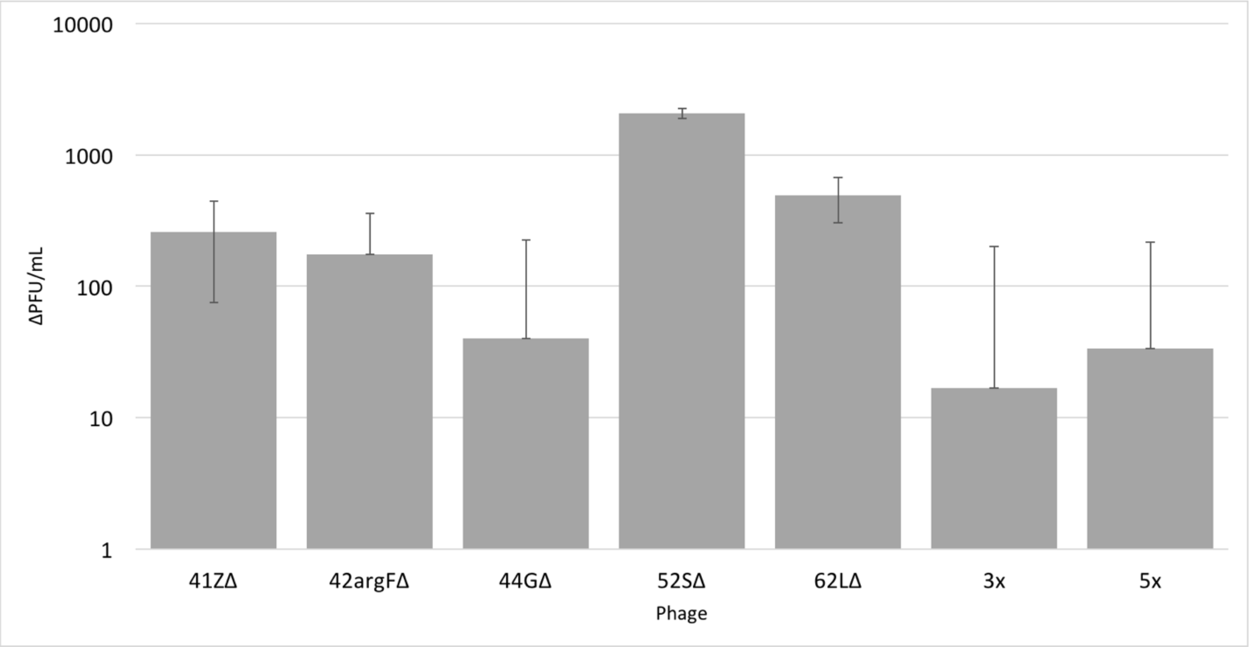


**Supplemental Figure 7. Phage titer increase over time during PAO1 infection.** The increase in phage titer from zero to four hour in liquid culture infection. Most phages increased modestly; two (41ZΔ*int*, and 52SΔ*int*) had a significant increase in phage titer. The phage cocktails had a negligible titer increase, which could suggest interactions between the phages in each cocktail. The Y-axis is in logarithmic scale, error bars are the standard deviation for all samples, n=3.

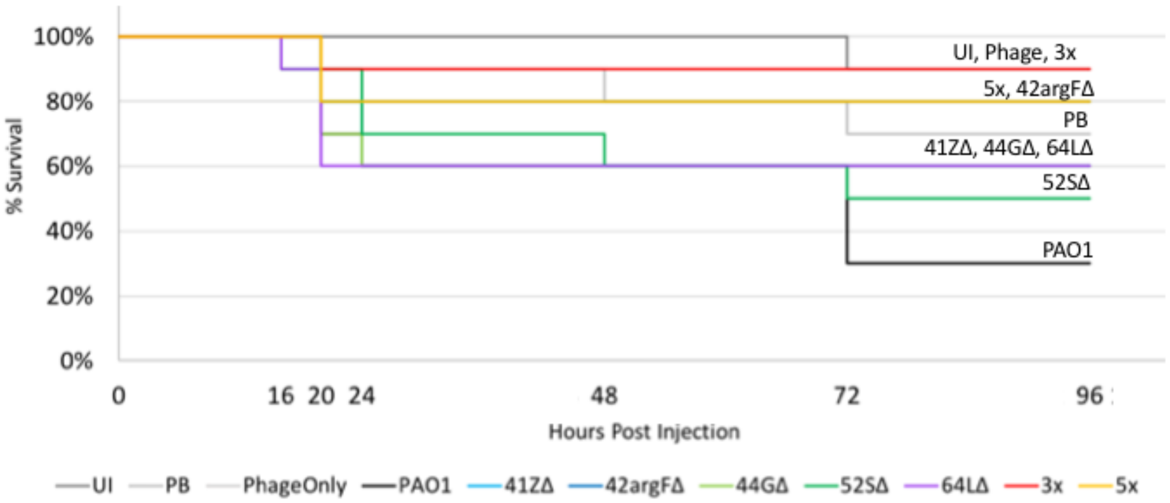


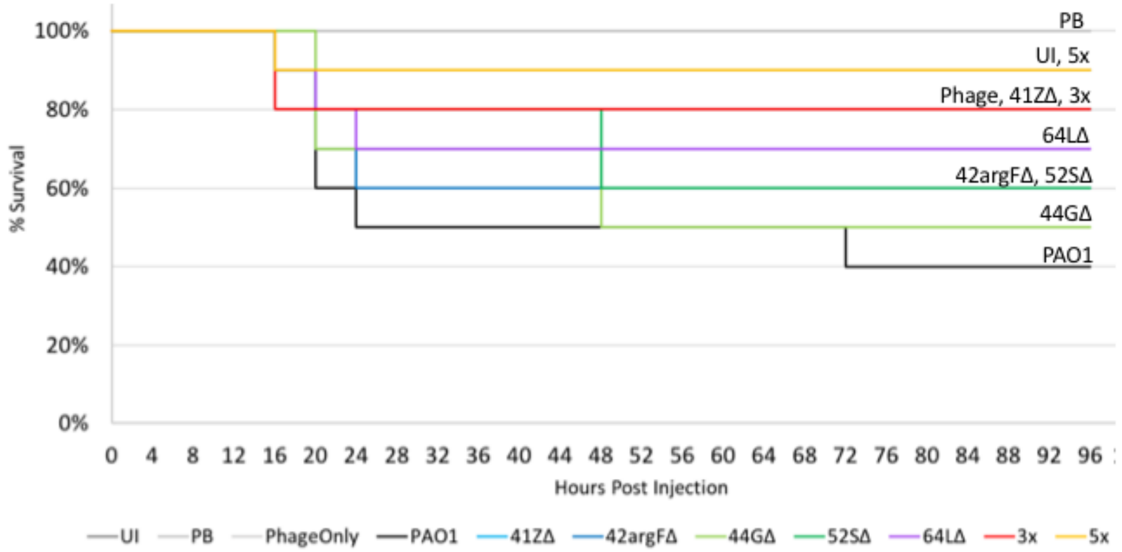


**Supplemental Figure 8. Engineered phages protect *Galleria mellonella* larvae from bacterial infection.** Two additional independent replicates of the experiment in Figure 4.

**Supplemental Tables**

**Supplemental Table 1. Integrative genetic element predictions for Pae5 and Pae1505**
Phages are highlighted in green.
Supplied as Excel

**Supplemental Table 2. Primers Used in Experiments**

| Primer Name | Sequence (5' -> 3') | Use |
|---|---|---|
| Pae5.52yheS_L | cacgccgcgctggatgct | Detection of CJ for 52yheS WT phage |
| Pae5.52yheS_R | acggctcggagagcagct | |
| Pae5.42argF_L | gtccgacagcacaacgag | Detection of CJ for 42argF WT phage |
| Pae5.42argF_R | cggcaatcctgcctgccg | |
| Pae5.64L_L | cctatctctatttccgcc | Detection of CJ for 64L WT phage |
| Pae5.64L_R | aaagcggaagtgccagct | |
| Pae5.41Z_L | aaacactgtgggtttacg | Detection of CJ for 41Z WT phage |
| Pae5.41Z_R | cgaggcgacacgatttcg | |
| Pae1505.43spxA_L | cataagcgccgcgttcgc | Detection of CJ for 43spxA WT phage |
| Pae1505.43spxA_R | cgcggccatccgtccggc | |
| Pae1505.52S_L | gcatcgcggcgagccatt | Detection of CJ for 52S WT phage |
| Pae1505.52S_R | atccgcccgaaagcagcc | |
| 141.44G.R | CGCAGTCTTGAATTCGGGCG | Detection of CJ for 44G WT phage |
| 142.44G.L | GCCGTTCTCCCGCACCTTTA | |
| 65.52SL.dx | AGCACGCCGATGGACAGAT | Detection of DJ for 52S Δ*int* phage |
| 67.52SR.dx | GGCGGAGGTATGTTATCCCG | |
| 88.Pae5.41Z.dxL | GTCGAAGGGCGGCAAGAAAG | Detection of DJ for 41Z Δ*int* phage |
| 89.Pae5.41Z.dxR | GCGACTACACAACCGTCTCA | |
| 96.Pae5.42argF.dxL | TCGGCAGATAGGCAGTTCCG | Detection of DJ for 42argF Δ*int* phage |
| 97.Pae5.42argF.dxR | AGTGTGAGCCAGACGTGCTT | |
| 133.52yheS.dtA | CAGGAAGGAAGGAGGTGGGG | Detection of DJ for 52yheS Δ*int* phage |
| 134.52yheS.dtD | GTGCAGCGATTTTCGGCAGG | |
| 157.44G.dxL | CTCAGGGAGGGCCACGCGAT | Detection of DJ for 44G Δ*int* phage |
| 158.44G.dxR | GCTGCGAGATGAGTCGCGTG | |
| 163.64L_dxRnew | GGAAATACTCACTTCCCGGC | Detection of DJ for 64L Δ*int* phage |
| 165.44G_dxRnew | GCTGCGAGATGAGTCGCGTG | |

**Supplemental Table 3. Primers used for phage engineering** Primer name, sequence, length of product, and overlap are shown.

| Primer Name | Sequence (5' -> 3') | Length of Product (bp) | Overlap with Right Side (Fragment) |
|---|---|---|---|
| 52S.Af | GCGGCATCCAGAGAATGAGAAATCATCCGGGGTGGGAGAT | 12463 | 42bp (B) |
| 52S.Ar | GAGGAATCCCGCGAGTGGAA | | |
| 52S.Bf | GCTTCTTGACGACGCGGTAA | 12574 | 41bp (C) |
| 52S.Br | CTACGCCCGTTGGTGTCCTT | | |
| 52S.Cf | CAACGAGACGCACCCCGA | 12619 | 38bp (D) |
| 52S.Cr | TGATCCAATGAACGGTCAGCA | | |
| 52S.Df | TGTAAACGGCACGAATGCTG | 13121 | 38bp(C) |
| 52S.Dr | ATCTCCCACCCCGGATGATTTCTCATTCTCTGGATGCCGC | | |
| 41Z.Af | TGATCTGCCGAGGTGAAAGCCGCGTCTTCGGTGTAGCCAGA | 14325 | 39bp (B) |
| 41Z.Ar | GCGGAGACGGAATGCCTTTG | | |
| 41Z.Bf | TATCCGTCGCATGGCCTGTT | 11936 | 48bp (C) |
| 41Z.Br | AGTGCCCTCCAAGGATGACC | | |
| 41Z.Cf | TGCAGTGTATTCCGTCGCTCA | 13279 | 41bp (A) |
| 41Z.Cr | TCTGGCTACACCGAAGACGCGGCTTTCACCTCGGCAGATCA | | |
| 42argF.Af | GCAATGGGGCTGCTCGTTCATATAACCCCGCACAACCCC | 13515 | 39bp (B) |
| 42argF.Ar | CCTACAAGTCTGCCACCGTC | | |
| 42argF.Bf | GTCAGCGCCTCGATCACATC | 13025 | 45bp (C) |
| 42argF.Br | TGTGGAGCGTACTCAACGAC | | |
| 42argF.Cf | GGCTGGGTATCGCTAACCATTA | 13559 | 39bp (A) |
| 42argF.Cr | GGGGTTGTGCGGGGTTATATGAACGA GCAGCCCCATTGC | | |
| 44G.Af | GAGCGGGCGGAGGGAATCGACCCACGCGGCCGGGAGAGAG | 14317 | 47bp(B) |
| 44G.Ar | ACCTCGCCGGAGCAACCACA | | |
| 44G.Bf | GCGGCGCTGCTGGTGGTGAC | 13561 | 42bp(C) |
| 44G.Br | TGCGCTGGTCCGAGAGCGAT | | |
| 44G.Cf | GAGTCCTCGGACAAGGCGGCCAA | 14426 | 40bp (A) |
| 44G.Cr | CTCTCTCCCGGCCGCGTGGGTCGATTCCCTCCGCCCGCTC | | |
| 52yheS.Af | ATGGTGCCTGAAAGCCGTGATTTCTGCTTGTTGCGGCGAG | 12648 | 52bp (B) |
| 52yheS.Ar | ATCACGGCGAAGGTAGACGG | | |
| 52yheS.Bf | GCGTCGGTCAATTCGTGCTT | 12376 | 46bp (C) |
| 52yheS.Br | GAGCTCGTCCACCAGGGAT | | |

| | | | |
|---|---|---|---|
| 52yheS.Cf | AGACCATCGTTGGCTTCAAGGT | 12641 | 45bp (D) |
| 52yheS.Cr | TCGATGTTCTTGCGTGCGACC | | |
| 52yheS.Df | CAGCCGAGGAGGAGACCAAG | 12561 | 40bp (A) |
| 52yheS.Dr | CTCGCCGCAACAAGCAGAAATCACGGCTTTCAGGCACCAT | | |
| 64L.Af | GGATGGCGCATTTGAAGGCATCACTCCGCGGCCTCTTCTTC | 13672 | 49bp (B) |
| 64L.Ar | GGTCACCGTAGGAGCATCCG | | |
| 64L.Bf | CGAGCCTTGGACTACGTGTG | 15858 | 47bp (C) |
| 64L.Br | GGTGGTTTTGCTTGCGGTCA | | |
| 64L.Cf | AACCGGAACATGTAAGGGAGC | 15751 | 41bp (D1) |
| 64L.Cr | CTGCAGGGTGGCCTTGTAGG | | |
| 64L.D1f | AAGTTCTTGGCGGACGTGGA | 7999 | 47bp (D2) |
| 64L_D1r | GCGGCTGACGGACAGAAGAT | | |
| 64L_D2f | GAGGCCTACACGGACTGGG | 8907 | 41bp (A) |
| 64L.D2r | GAAGAAGAGGCCGCGGAGTGATATCGCGGCAGCTAATGCTT | | |
